# Cancer Classification by Correntropy-Based Sparse Compact Incremental Learning Machine

**DOI:** 10.1101/028720

**Authors:** Mojtaba Nayyeri, Hossein Sharifi Noghabi

**Affiliations:** Department of Computer Engineering, Ferdowsi University of Mashhad, Iran; Center of Excellence on Soft Computing and Intelligent Information Processing.

## Abstract

Cancer prediction is of great importance and significance and it is crucial to provide researchers and scientists with novel, accurate and robust computational tools for this issue. Recent technologies such as Microarray and Next Generation Sequencing have paved the way for computational methods and techniques to play critical roles in this regard. Many important problems in cell biology require the dense nonlinear interactions between functional modules to be considered. The importance of computer simulation in understanding cellular processes is now widely accepted, and a variety of simulation algorithms useful for studying certain subsystems have been designed. In this article, a Sparse Compact Incremental Learning Machine (SCILM) is proposed for cancer classification problem on microarray gene expression data which take advantage of Correntropy cost that makes it robust against diverse noises and outliers. Moreover, since SCILM uses l_1_-norm of the weights, it has sparseness which can be applied for gene selection purposes as well. Finally, due to compact structure, the proposed method is capable of performing classification tasks in all of the cases with only one neuron in its hidden layer. The experimental analysis is performed on 26 well known microarray datasets regarding diverse kinds of cancers and the results show that the proposed method not only achieved significantly high accuracy but also because of its sparseness, final connectivity weights determined the value and effectivity of each gene regarding the corresponding cancer.

## 1. Introduction

Most of human diseases are influenced by genes, and identifying genetic landscape and profile of diseases is an undisputable fact especially when it comes to diseases such as cancer [1]. In the quest for determination of genetic causes of diseases, new technologies such as Next Generation Sequencing [2, 3] or Microarray expression [4] which are high-throughput procedures have paved the way to quantitate and record thousands of genes expression levels simultaneously [5-7]. These new technologies provide computational oncologists with valuable information for cancer prediction and cancer classification [1, 8, 9]. Making the best use of these valuable information and extracting it from datasets requires advanced, accurate and robust computational techniques because these datasets most of the time follow “large-p-small-n” paradigm which means they have high number of observed genes but low number of samples [10]. Cancer classification has been studied comprehensively with diverse methods from weighted voting scheme [11] and Partial Least Square (PLS) [6] to Support Vector Machines (SVM) [12] and Extreme Learning Machines (ELM) [13]. In addition to these methods, Artificial Neural Networks (ANNs) [14], Probabilistic neural networks (PNNs) [15] and soft computing approaches (hybrid of evolutionary computation and machine learning) were also applied and developed for cancer diagnosis and cancer classification [10, 16]. One of the well-known types of ANNs are constructive networks whose optimum structures (number of nodes in the hidden layer) are determined automatically [17-19]. In these networks, number of nodes and connectivity weights are gradually increased from the lowest to the optimum value and they are categorized in two types: compact [17, 18, 20] and non-compact [19]. Input parameters of the newly added node in the non-compact type are specified randomly whereas in the compact one, they are adjusted via an optimization process.

Most of these methods are suffering from “curse of dimensionality” which is related to high dimensions of these datasets. Another aspect of cancer classification is related to feature selection (gene selection) methods in order to prevent over-fitting in the learning process [21]. Model *et al.* [22] applied several feature selection methods for DNA methylation based cancer classification. Another comparative study for feature selection was performed by Li *et al.* [23] for tissue classification based on gene expression. Cawley *et al.* [24] proposed a sparse logistic regression with Bayesian regularization for gene selection in cancer classification and Zhang *et al.* [25] used SVM with non-convex penalty for the same problem. Piao *et al.* [26] take advantage of ensemble an correlation-based gene selection method for gene expression data regarding cancer classification. Interested readers can refer to five good surveys of feature selection in [27-30] and [31] and the references therein. However, feature selection comes with certain prices such as addition of another layer of complexity to the model or information loss [27].

In this article, we propose Sparse Compact Incremental Learning Machine (SCILM) which prevents over-fitting without feature selection due to its compact structure. Further, because of Correntropy cost SCILM is robust against noises and outliers. In addition to these advantages, since SCILM takes advantage of l_1_-norm of the weights, it is sparse and this sparseness determines the most effective connectivity weights corresponding to all features. Therefore, the final weights of the generated model by SCILM can be utilized for gene selection purposes as well. SCILM is a learning method for datasets with low sample size and high dimensions. These characteristics are highly important and medical and pharmaceutical research because numbers of genes or drug compounds are significantly lower than number of features and attributes one can find for them. SCILM is proposed for such problems and microarray profiles for cancer classification have both these characteristics. The presented method prevents over-fitting without feature selection due to its compact structure and also because of Correntropy cost SCILM is robust against noises and outliers. Authors in [32], investigated robustness of Correntropy objective function. In addition to these advantages, since SCILM takes advantage of l_1_-norm of the weights, it is sparse and this sparseness determines the most effective connectivity weights corresponding to all features. Therefore, the final weights of the generated model by SCILM can be utilized for gene selection purposes as well.

The rest of the paper is organized as follows: section 2 presents the proposed method, section 3 describes the results and final section concludes the paper.

## 2. Methods and Materials

This section presents a new constructive network with sparse input side connections. The network has a single hidden layer in which the hidden nodes are added one by one until the network reaches a certain predefined performance. After the new hidden node is added and trained, its parameters are fixed and do not changed during training the next nodes. Each newly added node is trained in two phases: a) Input parameters adjustment, b) output parameter adjustment. The input parameters of the newly added node are trained based on Correntropy objective function. The output connection is adjusted by MSE objective function. In the rest of this section some preliminaries are described followed by the description of the proposed algorithm.

### 2.1 Dataset representation

The dataset with *N* distinct samples is denoted by

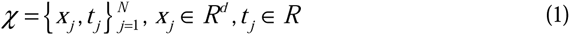

### 2.2 Network structure

Let f be a continuous mapping, f_L_ be the output of the network with L hidden nodes. The network is represented as

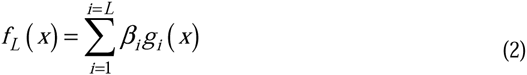

Where

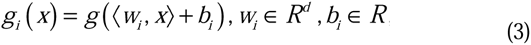

Where 〈.,.〉 is inner product between two elements. **In this paper g is considered as tangent hyperbolic function**. The network (with L hidden nodes) error vector is defined as

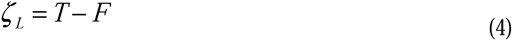

Where *T* = [*t*_1_, …, *t_N_*] and *F* = [*f_L_*(*x*_1_), …, *f_L_*(*x_N_*)] The activation vector for the *i*th hidden node is

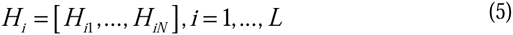

where *H_ij_* = *g_i_*(*x_j_*), *j* = 1, …, *N*;*i* = 1, …,*L*.

### 2.3 Correntropy

Let v and u be two random variables with *ζ* = *u* − *v*. The Correntropy is a similarity measure between two random variables and defined as

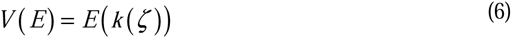

Where *E*(.) denotes the expectation in probability theory and k(.) denotes a kernel function which satisfy Mercer condition. In this paper only the Gaussian kernel is used. Regard to this,

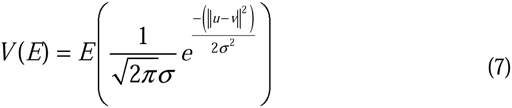

### 2.4 Proposed method

This subsection proposes a new incremental constructive network with sparse hidden layer connections. The hidden nodes are added to the network and trained one by one. When the new node parameters are tuned, they are frozen and do not change during training the next nodes. Fig. 1 illustrates the mechanism of the proposed method.

**Fig. 1.**
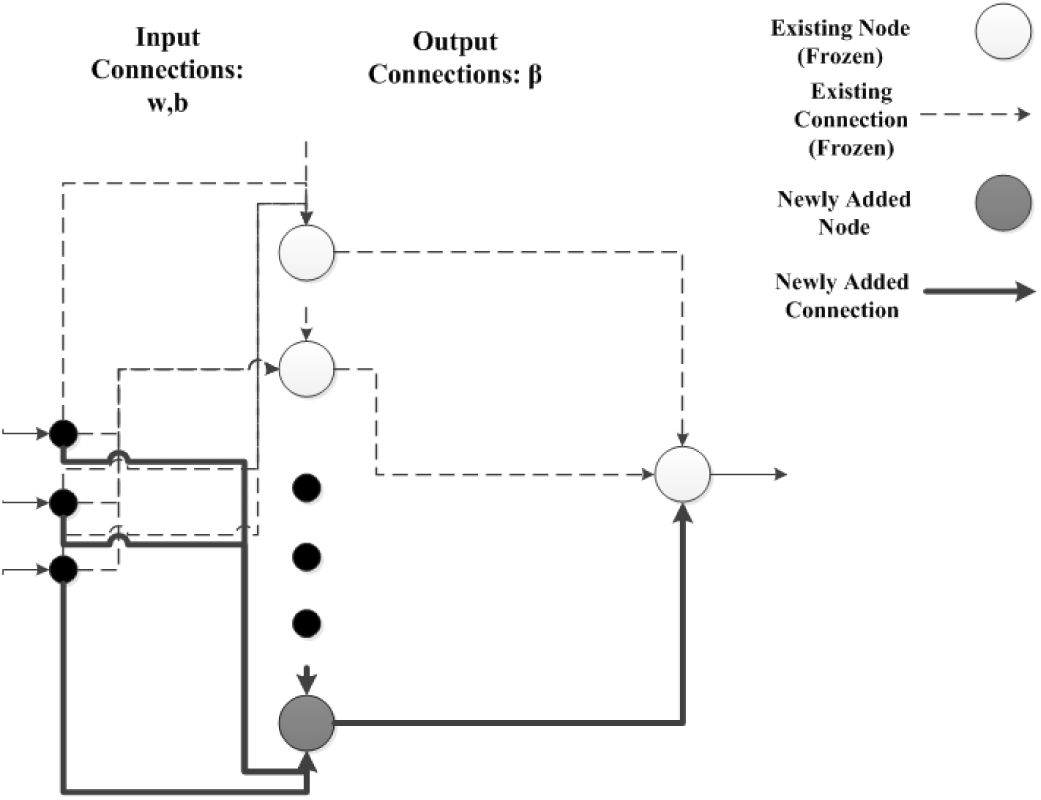
Each newly added node is trained in two stages: a) Input weights adjustment b) Output weight adjustment. Existing nodes and connections do not change during training the new node. (R1.1), (R1.2), (R2.2)

**Fig 1a.**
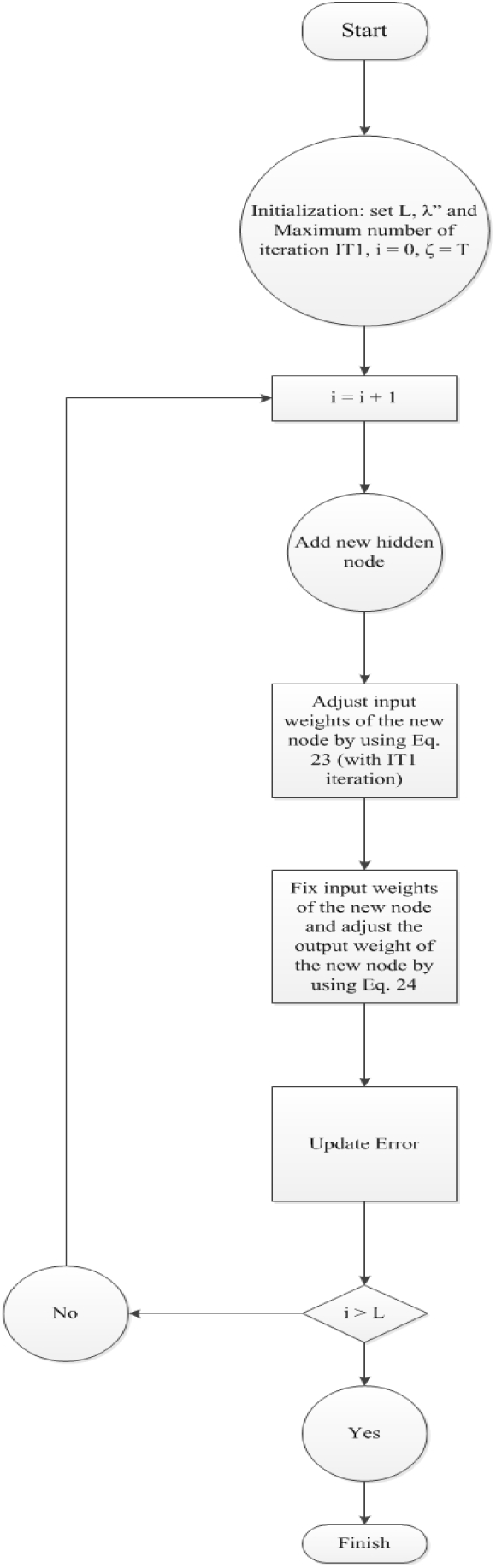
Flow chart of the proposed algorithm. (R1.2)

Training of the new node performs in two stages:

#### Stage 1: input side optimization

In the previous work [33], Input parameters of the new node are trained based on Correntropy objective function as follows:

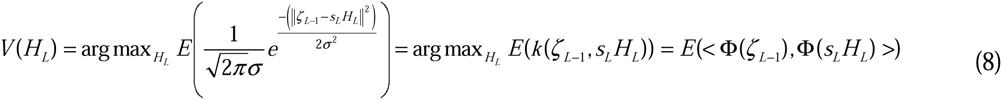

Where *s_L_* is a real number which is obtained by trial and error and *ζ_L_*_–1_ is the residual error for the network with *L*-1 hidden nodes. Regarding Eq.(8), the new node *H_L_* has most similarity to the residual error (regard to kernel definition). It is important to note that when the new node vector equals to the residual error vector (most similarity between the new node and the residual error), the training error becomes zero. Thus the optimal condition is [33]

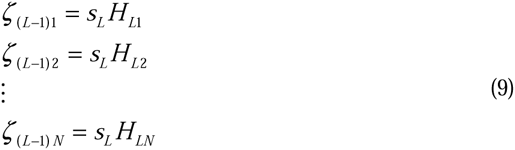

Similarly, it is known that *H_Li_* = *g_L_* (*x_i_*), *i* = 1,…,*N* and *g* is tanh(.). Since *g* is bipolar and invertible the system (9) can be rewritten as [33]

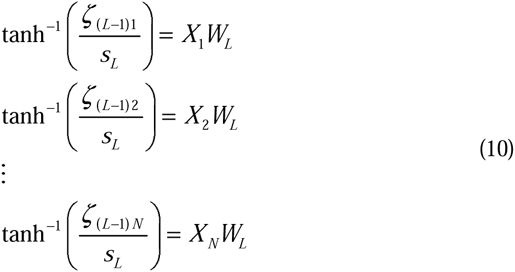

Where 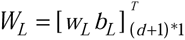, wL = [*w_L_*_1_,…,*w_Ld_*]_(_*_d_*_*1)_, *X_i_* = [*x_i_* 1]_1*(_*_d_*_*1)_, *i* = 1,…,*N* and *w_L_, b_L_* are input connections (input weight and bias) of the *L*th hidden node and *x_i_* is the i*th* training sample. To obtain added simple representation, let:

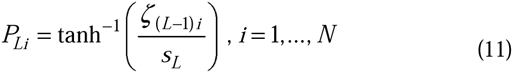

Thus, we can write

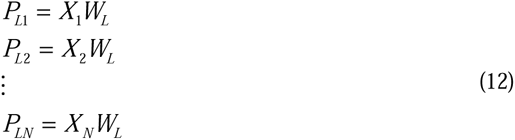

Regard to this, as mentioned in [33], and since system of equations (9) and (12) are equivalent, the following equation will be solved instead of (8):

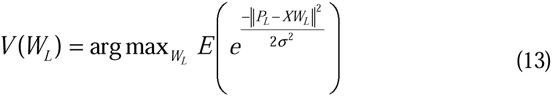

Where

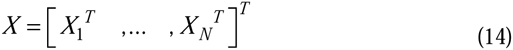

The expectation can be approximated from data points, the constant term can be removed, and thus the following optimization problem is obtained [33]:

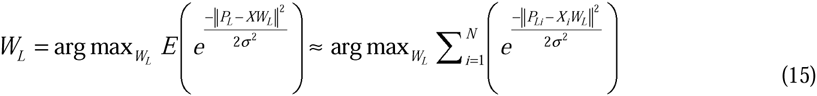

As mentioned in [33], to avoid overfitting and achieve a better generalization performance, the regularization term should be added:

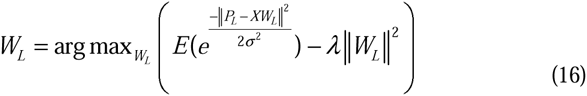

As mentioned in [33] and similar to [34], employing the half quadratic optimization problem, the local solution of (16) is obtained, using the following iterative process [33]:

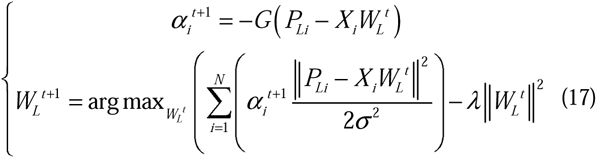

Where 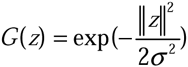. It is obvious that auxiliary variables i.e., 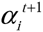 help to reduce effect of data that are contaminated by noises. After some derivations (multiplying the optimization problem by constant term *σ*^2^ and set *λ*′ = 2*λ*) which are mentioned in [33], we obtain:

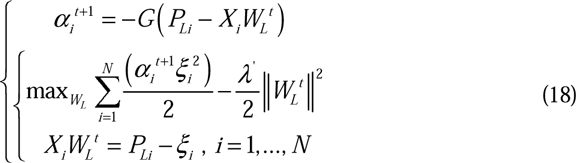

Several literatures in machine learning and adaptive filters replaced l2 norm by l1 norm to provide sparse solution (optimum weight) [35, 36]. Accordingly, inspired by [36] and Different from previous work [33], to provide a sparse solution, the following iterative process should be performed instead of (18):

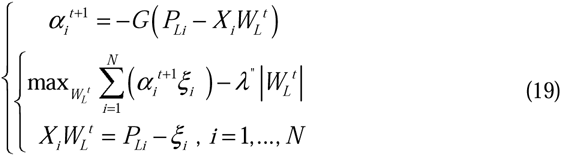

Where|.| denotes the *l*_1_ norm. Consider the following optimization problem that is extracted from Eq. (19):

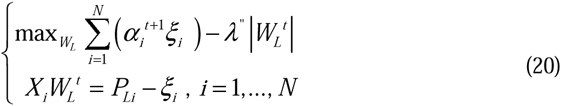

Let *r* = [*r*_1_,…*r_d_*_+1_], *s* = [*s*_1_,…,*s_d_*_+1_], *p* = [*p*_1_,…,*p_N_*] and *q* = [*q*_1_,…,*q_N_*], where r, s ≥ 0 and p, q ≥ 0.

Let 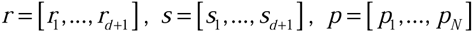 and *ξ_i_* = *p_i_* – *q_i_*, *i* = 1,…,*N*, inspired by [36], the optimization problem (20) can be rewritten as a linear programming problem:

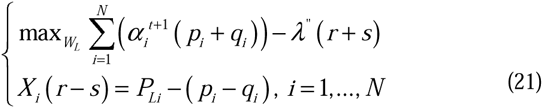

In order to solve the problem (21) by using Matlab toolbox, it needs to change optimization problem (21) to the standard form. The standard form of linear programming which is used in Matlab is:

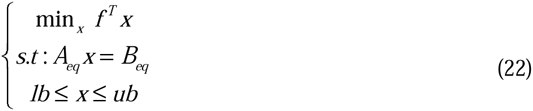

Thus, passing from (21) to (22), we obtain:

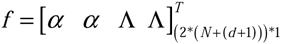

Where

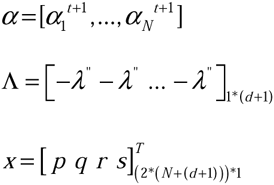

And

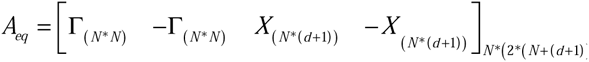

where

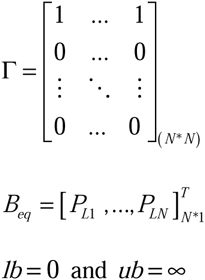

Thus, the optimum input parameters of the new node are adjusted using the following iterative process:

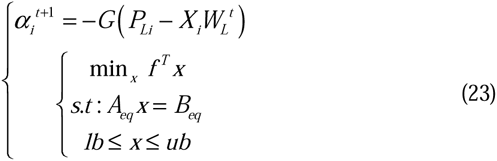

#### Stage 2: Output side optimization

Similar to [17], the output weight is adjusted using the following equation

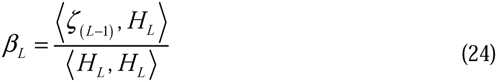

Where *H_L_* is obtained from previous stage.

The proposed method is specified in Algorithm 1 and Fig. 2 is the flow chart of SCILM:

**Fig. 2.**
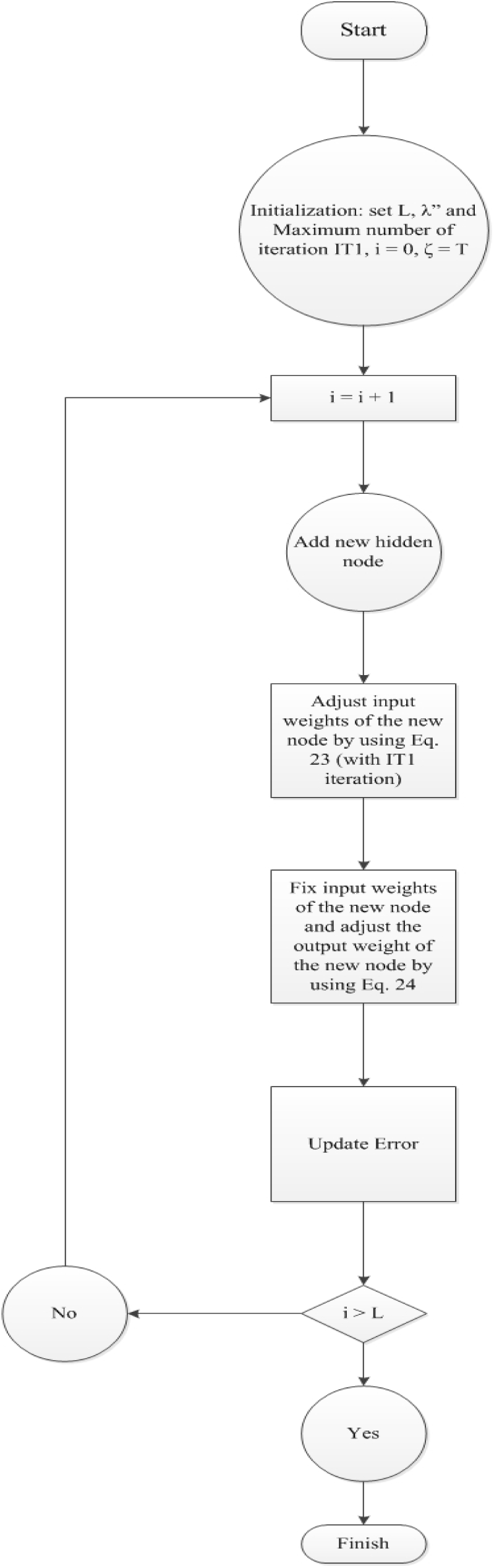
Flow chart of the algorithm. (R1.2)

##### Algorithm 1 SCILM

**Input:** training samples 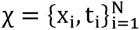

**Output:** The optimal input and output weights β_i_, W_i_, i = 1,…, L

**Initialization:** Maximum number of hidden nodes *L*, regularization term *λ*″, Maximum number of iterations IT1

**For** i=1:L

Stage 1: calculate P_L_ and X by (11) and (14)

For k=1:IT1

Update input parameters according to (23)

END

Stage 2: calculate the hidden node vector H_L_ using (5), and the error vector, *ζ_L_*_−1_, using (4):

Adjust the output weight according to (24)

END

Update Error as *ζ_L_* = *ζ_L_*_–1_ − *β_L_ H_L_*

**END**

### 2.5 Datasets

The experiments were performed on 26 datasets adopted from [21] which contain 11 multi-class and 15 two-class datasets. We applied the same reference numbering system as [21] for convenience. Reference numbers less than or equal to 15 are for two-class datasets and reference numbers greater than 15 indicate multi-class datasets. Most of the datasets have dimensions in the range of approximately 2000 to 25000 (except dataset 19 about yeast which has 79 features) and the sample size varies approximately from 50 to 300. The detail for each dataset is as follow:

Six of the datasets have been studied in [37] and among them, datasets numbered 1 to 3 relate to breast cancer, 4 and 5 deal with lung cancer and dataset 6 is about hepatocellular carcinoma. Another group of six datasets studied in [38]. Datasets 7 and 8 deal with prostate cancer, 9 and 10 are about breast cancer and finally 16 and 17 are related to leukaemia. Five well-known bioinformatics datasets are about colon cancer (11) [39], ovarian cancer (12) [40], leukaemia (13) [11], lymphoma (18) [41] and yeast (19) [42]. There rest of the datasets are selected from NCBI GEO and their corresponding IDs are available in [21] and in this paper they are numbered in the range of {14,15} U {20,&,26}.

### 2.6 Experiments

This paper used SVMs with Radial Basis Function (RBF) and sigmoid kernels and Correntropy based ELM (ELMRCC) [34] to compare with SCILM. The RBF kernel used in SVM is defined as K(u,v) = exp(–γ||u – v||^2^) and the sigmoid kernel is defined as *K*(*u,v*) = tanh(*α*.*u.v* + *β*) K(u, v) =tnh(αuv + β). For SVM, we used the LIBSVM toolbox [43], the regularization parameter C is selected from the set {10^−3^, 10^−2^, …,10^5^} and the RBF kernel parameter γ is selected from{{10^−2^,…,10^2^}}. For the SVM with sigmoid kernel, the parameters α and β are selected from the sets {{10^−2^,…,10^2^}} and {10^−1^,…,10^1^} respectively. ELM-RCC [34] have a hyper parameter λ and the best of this parameter is selected from the set{{10^−3^,10^−2^,…,10^5^}}. Similar to [44], the number of hidden units in ELM-RCC is set to 1000. This paper used additive nodes with sine activation function for ELM-RCC as *h*(*x*)sin(〈*w*,*x*〉 + *b*) h(x) = sin(< *w*, *x* > + *b*). The parameters w, b are randomly generated by uniform distribution between -1 and 1. Furthermore, the data samples are normalized into range -1 and 1. For the proposed method, the parameter λ″ in the optimization problem (20) is selected from the set {10^−3^, 10^−2^, …, 10^5^} and the number of hidden nodes is set to one (the most compact network i.e., one hidden layer network with one hidden node) and the kernel width (*σ*) is selected from{10^−2^, …, 10^2^}. SVM used one against all strategy for multiclass classification datasets. To evaluate the performance of the proposed method in comparison with SVM and ELM-RCC, the testing accuracies are reported in table 1 and table 2. For the proposed method experiments are performed in 20 independent trials for each problem. In each trial, data samples are reshuffled and mean of accuracy is reported in the following tables.

## 3. Result and Discussion

In this section, we discuss the results from two viewpoints of accuracy and feature selection. In the accuracy part in comparison with the stated methods, SCILM achieved significantly better results in 10 datasets, performed equally in 6 datasets and only lost in 7 datasets. In table 1, the average accuracy of each two-class dataset over 20 independent runs is reported. According to this table, in the case of dataset 1 SCILM achieved the accuracy of 88.75% and among the compared methods ELM-RCC achieved 76.875%. For dataset 2, SCILM has an accuracy of 71.58% while SVM with RBF kernel has 66.94%. In the third datasets we achieved 71.33% but both SVMs were not able to perform better than 64.21%. In dataset 4, SCILM and SVM with RBF had accuracy levels of 61.27% and 58.09%, respectively. For the fifth dataset, SVM with sigmoid kernel and SCILM performed equally, however, in dataset 6 the same SVM performs better than the proposed method. Both SVMs achieved almost the same result as SCILM for dataset 7 in spite of the fact that SCILM was slightly better. In dataset 8, SCILM achieved 97.85% and ELM-RCC was not better than 94.28%. In dataset 9, SCILM shows the performance of 87.5% while SVM with RBF had the accuracy of 80%. In datasets 10, 12, 13 and 15 the proposed methods and the best of the compared methods achieved almost the same accuracy level; however, in datasets 11 and 14 SVM with RBF kernel performs better than the proposed method.

The results of the average test accuracies for multi-class datasets are reported in table 2. According to this table, for datasets 16 and 17 the proposed method achieved significantly better results than the best of the other compared methods, however, in datasets 18 and 19 SVM with sigmoid kernel achieved slightly better results than SCILM and for datasets 20 and 21, ELM-RCC achieved higher accuracy than SCILM. For the rest of the datasets, SCILM obtained competitive results and outperforms other methods especially in the case of datasets 22, 23 and 25.

It is important to note than SCILM has only 1 neuron in its hidden layer for all datasets, while ELM-RCC has 1000 neurons in its hidden layer and SVMs also take advantage of cross validation for parameter optimization based on LIBSVM library.

Concerning the aspect of feature selection, because SCILM is taking advantage of the l_1_-norm, it has sparseness and the final connectivity weights also tend to be sparse. Therefore, after generating the model for each dataset these final weights can be analyzed from the feature selection point of view. According to this approach, each feature has a corresponding weight which indicates the value of that feature i.e. more valuable features have higher values for their corresponding connectivity weights and less vital features have approximately equal to zero values for their final weights. Due to sparsity of the generated model, most of these weights tend to be near or equal to zero. The values of these weights for 8 randomly selected datasets are illustrated in figure 3. In order to save space the rest of the datasets are not considered in this figure. As shown in this figure, the weights are clearly specifying valuable features and separate them from less informative ones. The advantage of this feature selection approach is that it does not have additional computational cost or complexity for the model since it is within the learning process.

**Fig. 3.**
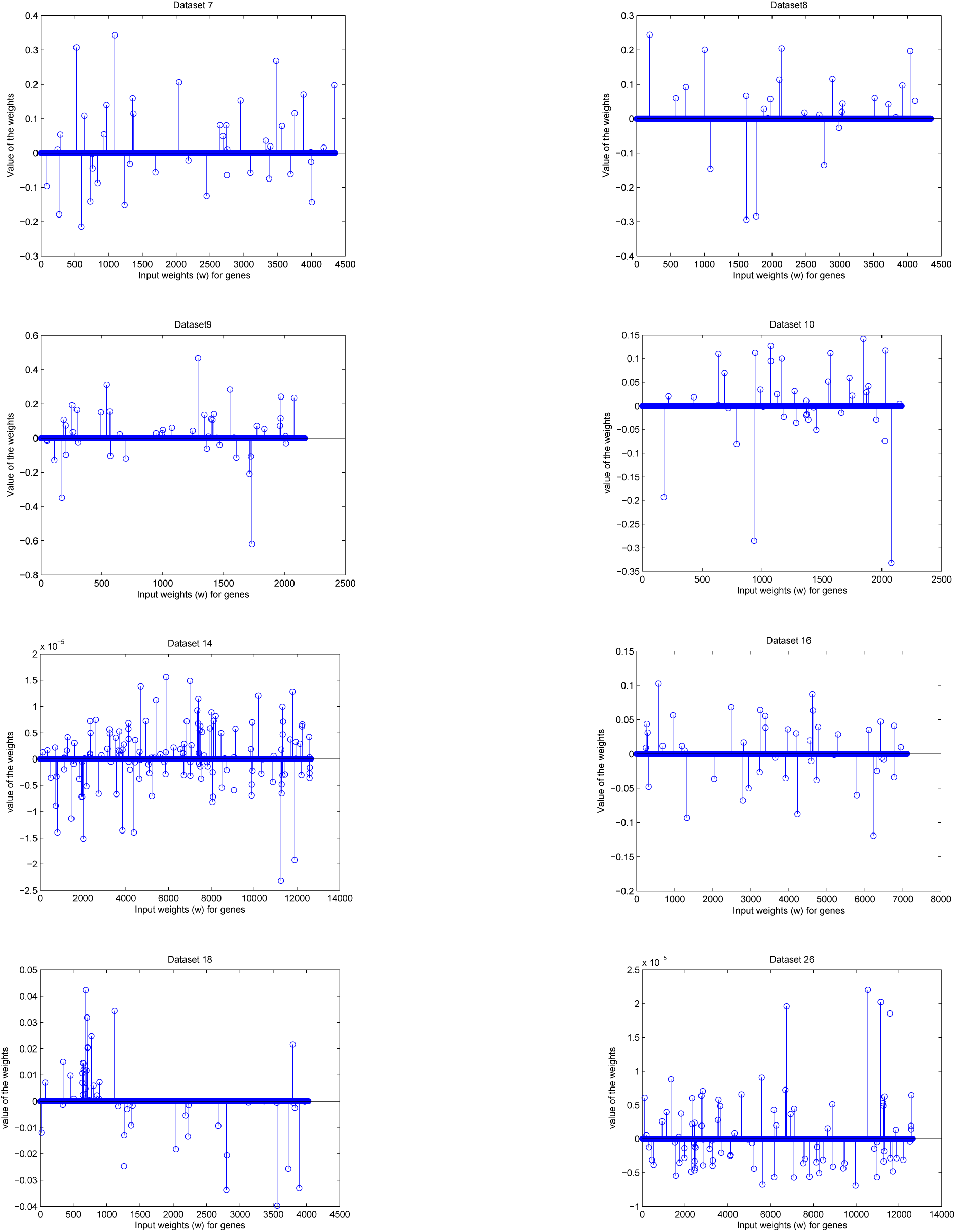
Connectivity weights (w) of the hidden layer are sparse and as these weights are coefficients of the genes, hardly informative genes are eliminated and only effective genes are considered for the model.

As for another advantage of SCILM, since the auxiliary variables (*α*) that appeared in optimization problem (20), have a small value for the outliers and these data points have a small role in the optimization of the network parameters, the proposed method is also robust to outliers as well.

The proposed network has an advantage of having a more compact architecture i.e., only has one hidden node in single layer, while the experiments demonstrated that SVM and ELM have several nodes (from 15 to 200 support vectors for SVM and 1000 hidden nodes for ELM). Furthermore, SCILM has the most sparse input side connection (in most cases sparsity rate is up to 99%). This simpler structure and the lowest number of degrees of freedom (parameters) lead to a substantially better generalization performance compared with the other methods.

## 4. Conclusion

In this paper, we have proposed a new classification method named SCILM based on incremental learning machines for cancer classification based on gene expression microarray data, which has two main significant advantages. First, because of Correntropy cost function, it is robust against uncertainty and noisy data. Second, since it uses the l_1_-norm of the weights in its optimization process, it can be seen as a feature selection method as well. This norm provides the generated model with sparseness in the weights, which can be exploited for feature selection purposes.

In the proposed method, the network structure is determined automatically, leading to a better generalization performance. Furthermore, due to optimization of the hidden layer parameters, the network has a compact architecture and fast training and testing speed.

We demonstrated that SCILM significantly improved the performance of other compared methods in both two-class and multi-class datasets. The capability of the SCILM for feature selection and selecting meaningful genes still requires more experiments and studies which is in center of our future research.

